# Examining the heritability of functional brain networks in adolescence

**DOI:** 10.1101/2025.08.08.669358

**Authors:** Christian Coffman, Sanju Koirala, Robert Hermosillo, Jacob Lundquist, Gracie Grimsurd, Oscar-Miranda Dominguez, Kimberly B. Weldon, Michael Anderson, Thomas Madison, Steve Nelson, Jed T. Elison, Sylia Wilson, Damien A. Fair, Brenden Tervo Clemmens, Saonli Basu, Eric Feczko

## Abstract

Discovering mechanisms underlying mental illness requires disentangling genetic and environmental factors influencing mental health. Researchers have started investigating the brain’s role as a potential intermediate biomarker linking genes and environment to mental health; however, understanding how much genetics shapes adolescent brain function remains elusive. Using data from the ABCD study (5,247 unrelated, 330 dizygotic, 248 monozygotic twin subjects), we estimated the heritability of functional connectivity and topography using both SNP and twin data. SNP-based heritability was calculated using genetic correlation and the recently developed AdjHE-RE estimator. We found low SNP heritability for brain functional connectivity (median = 2e-10% Gordon, 5.8e-7% probabilistic) and topography (max = 2%). Twin estimates using ACE models replicated prior findings from the literature (median = 8.9e-6% Gordon, 6% probabilistic, 27% topography). This suggest that additive genetic effects are minimally associated with functional brain features in adolescents highlighting the importance of considering both genetic and environmental factors when studying the development of functional brain networks and their relevance to mental health.

## Introduction

The increasing prevalence of child and adolescent mental health disorders presents significant societal and personal challenges. Mental health care costs have increased over 45 percent since 2017 and now represent more than 47 percent of all pediatric health care costs (Loo et al., 2024). For individuals, these conditions can impair functioning, and even elevate suicidality, making it crucial to understand their underlying pathophysiology. Research has made substantial strides in uncovering genetic and environmental factors contributing to mental health outcomes, with polygenic risk scores (PRS, which aggregates effects across all SNPs as they relate to some other trait) and twin studies underscoring the role of genetics (Smoller, 2020). Simultaneously, environmental factors, including socioeconomic status (Xu et al., 2024), lead exposure (McFarland et al., 2025; Xu et al., 2024), neglect (Rutter et al., 2010), and even weather events ( Rataj et al., 2016; Smoller, 2020), have also been shown to impact mental health outcomes.

Measuring genetic and environmental contributions to mental health is important. Such delineation may help identify genetic predispositions as well as environmental risk factors to better personalize mental health disorder diagnosis and treatment. However, disentangling the effects of genes and environment on mental health remains challenging. Often, complex causal pathways complicate the relationship between genetics, epigenetics, environment, and mental health outcomes. Such complications obscures studies attempting to detect factors mechanistically relevant to or predictive of mental health outcomes (Holmes & Davey Smith, 2019). Thus, researchers have sought intermediate biomarkers to link genetic risk to maladaptive behavioral manifestations.

Recent neuroimaging studies have examined functional brain features as potential intermediate biomarkers linking genes and environment to mental health. Evidence suggests that genes linked to mental health conditions may also be associated with brain structure (Coffman et al., 2024), neurotransmitter systems (Cabana-Domínguez et al., 2022; Coffman et al., 2024)), and functional connectivity (FC) (Sun et al., 2025). More recently, another feature of brain organization-network topography has also been associated with psychopathology (Cui et al., 2022; Keller et al., 2023; Lynch et al., 2023). If brain function connects genetics and behavior, then understanding how much genetics influences brain function can show how genetics affects behavior.

Numerous studies have leveraged twin designs and behavioral polygenic risk scores (PRS) to estimate the association between genetics and the brain. Prior work using twin designs have found heritability of functional brain network connectivity to be somewhat heritable (Elliott et al., 2018; Miranda-Dominguez et al., 2018; Ribeiro et al., 2021; Xu et al., 2024). Additionally, twin studies have shown that functional network topography is also heritable (Anderson et al., 2021); (Adhikari et al., 2018). Furthermore, recent studies have examined polygenic risk and its association with functional connectivity for schizophrenia (Qi et al., 2022); (Cao et al., 2021), Alzheimer’s disease (Axelrud et al., 2019), autism spectrum disorder (Lawrence et al., 2022), ADHD (Hermosillo et al., 2020), and general psychopathology (Sun et al., 2025). Characterizing the association of genetics with functional brain features with twin and polygenic risk-score (PRS) based estimates is useful and can be further enhanced with the addition of SNP heritability estimates by directly associating genetics to the function brain features.

Many behavioral traits likely emerge from the interaction between highly correlated (Hart et al. 2021) genetic influences (risk) and external environmental factors (exposures) (Erdelyi-Hamza et al., 2025; Hart et al., 2021; Plomin et al., 1977). If a PRS is associated with exposure that is related to the outcome, it may misleadingly imply genetic associations with the outcome itself. For example, imagine a group of sailors with high genetic <u>risk</u> for tall height. Researchers might observe that high genetic risk is associated with concussions (outcome) which could be interpreted as genetic risk for concussions. However, in reality taller sailors are more likely to hit their heads on the low ceilings of ships (exposure), making concussions more frequent. The PRS for height is not relating biological risk to concussions, but rather the risk of exposure – hitting one’s head. Such an example poses complications for interpretation of PRS for: while one interpretation is that genes are associated with the brain that increases exposure (hitting one’s head), leading to the condition (concussions). Alternatively, genetics may be influencing an indirect trait (height) that increases exposure (hitting one’s head), which can lead to the condition (concussions) and brain changes. Directly measuring how much the brain is associated with genes clarifies how much of a PRS-brain correlation may be genetic. To our knowledge, genetic associations with adult functional brain organization are minimal ((Elliott et al., 2018)). However, genetic associations with adolescent functional brain organization have not been measured.

In this study, we elucidate the association of genetics to functional brain organization by directly estimating the heritability of FC and network surface area in a large sample of 9-10 year old adolescents from the Adolescent Brain and Cognitive Development (ABCD) study. Our estimates utilize the pairwise genetic correlation between subjects (the genetic relatedness matrix or GRM) in a variance component model to model contributions from additive genetic effects as opposed to individual variation. The proportion of the variance of the outcome explained by the GRM is called SNP heritability. We estimated SNP heritability using a recently established tool (AdjHE-RE) designed to control for batch effects which are common in multisite brain imaging studies such as the ABCD dataset. In addition, we replicate prior twin studies examining the heritability of functional connectivity and surface area. Then, we compare SNP to twin based estimates to provide additional context for the relative effects of additive genetic effects and common environmental effects.

## Results

We estimated SNP-based and twin-based heritability of functional brain organization. SNP estimates were based on a variance component approach, AdjHE-RE, that accounts for confounding from site/scanner effects (Coffman et al., 2024). Twin estimates leveraged the well documented ACE model (Wang et al., 2011). The study sample included 5,247 unrelated subjects for the SNP estimates and 330 dizygotic (DZ) twins, and 248 monozygotic (MZ) twins.

### Heritability of functional connectivity is minimal

We first estimated SNP-heritability for functional connectivity (FC) between brain networks defined via two parcellations: 1) Gordon parcellation (Gordon et al., 2016), which is based on group average maps and 2) MIDB probabilistic parcellations (Hermosillo et al., 2024), a newer parcellation method that adapts ROI boundaries based on a predefined consensus threshold across participants. This parcellation method is particularly useful to control for individual variability in network organization across participants. Across all ROI-ROI connections, SNP-based heritability estimates for FC were extremely low between Gordon ROIs (median = 2e-10%, IQR = 1%, Max = 39%) with 6% of total estimates reaching statistical significance after adjusting for multiple testing using Bonferroni’s method (Armstrong, 2014). The greatest proportion of heritable connections within or between two networks was Somatomotor Dorsal connections to the Dorsal Attention Network (SMD-DAN) where 0.1% of SMD-DAN connections were significantly heritable. Across all MIDB probabilistic FC: SNP-based heritability estimates were low (median = 5.8e-7%, IQR = 7e-5%, Max = 12%), with 6% being significant after Bonferroni correction with significance being distributed across multiple network-network connections (**Figure 2**).

**Figure 1:**
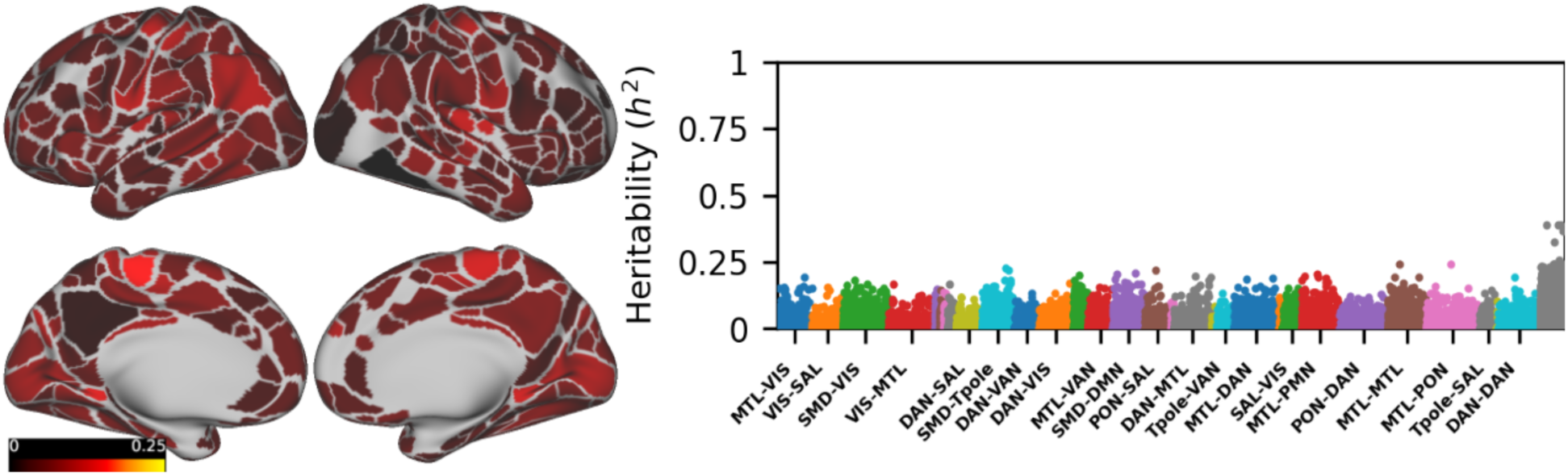
Left: ROI-wise projections of 90th percentile SNP heritability estimate for Fisher z-transformed functional connectivity values between Gordon parcels, mapped onto the Conte69 inflated cortical surface. The color scale spans 0 to 25% heritability for consistency across figures. Both lateral (top) and medial (bottom) views are shown for each hemisphere. Right: Manhattan-style plot of heritability estimates, with edges grouped and color-coded by the resting-state networks of the connected ROIs. The y-axis spans the full range of possible heritabilities (0-1). Network-network pairs accounting for fewer than 1.5% of total ROI-ROI connections are rescaled 100-fold on the x-axis and displayed in grey for clarity.

**Figure 2:**
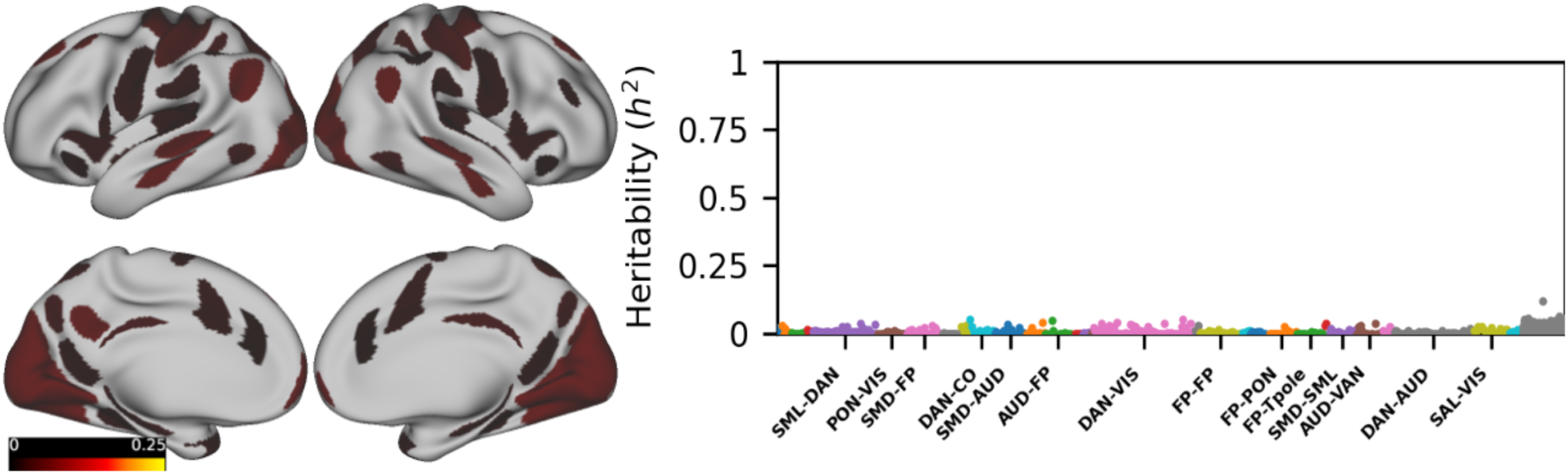
Left: ROI-wise projections of 90th percentile SNP heritability estimate for Fisher z-transformed functional connectivity values between MIDB probabilistic parcels, mapped onto the Conte69 inflated cortical surface. The color scale spans 0 to 25% heritability for consistency across figures. Both lateral (top) and medial (bottom) views are shown for each hemisphere. Right: Manhattan-style plot of heritability estimates, with edges grouped and color-coded by the resting-state networks of the connected ROIs. The y-axis spans the full range of possible heritabilities (0-1). Network-network pairs accounting for fewer than 1.5% of total ROI-ROI connections are rescaled 100-fold on the x-axis and displayed in grey for clarity.

We then estimated twin-heritability for functional connectivity (FC) using the same two parcellations. Twin-based heritability was slightly higher but still low overall between Gordon ROIs (median = 8.9e-6%, IQR = 3%, Max = 46%) with 25% of total estimates reaching statistical significance after adjusting for multiple testing using Bonferonni’s method. The largest proportion of network-network connections on the cortical surface were found in the Auditory to Dorsal Attention Network (AUD-DAN) with ∼4% of connections found significantly heritable (**Figure 3)**. Twin-based heritability estimates were higher but still relatively small between probabilistic ROIs (median = 6%, IQR = 7e-5%, max = 0.71%) with 28% being statistically significant after Bonferroni correction. The largest proportion of network-network connections found significantly heritable were observed in the Ventral Attention Network and Parietal Memory Network (VAN-PMN) with ∼30% showing significant heritability (**Figure 4**). The 5th and 95th percentiles for common environmental estimates spanned 0 to 25% for Gordon ROIs and 0 to 2% for MIDB probabilistic ROIs with correlations between variance components: genetic-common environment = –0.18 and –0.38, genetic-individual variation = –0.82 and –0.88, and common-individual variation = –0.40 and –0.08 for each parcelation, respectively.

**Figure 3:**
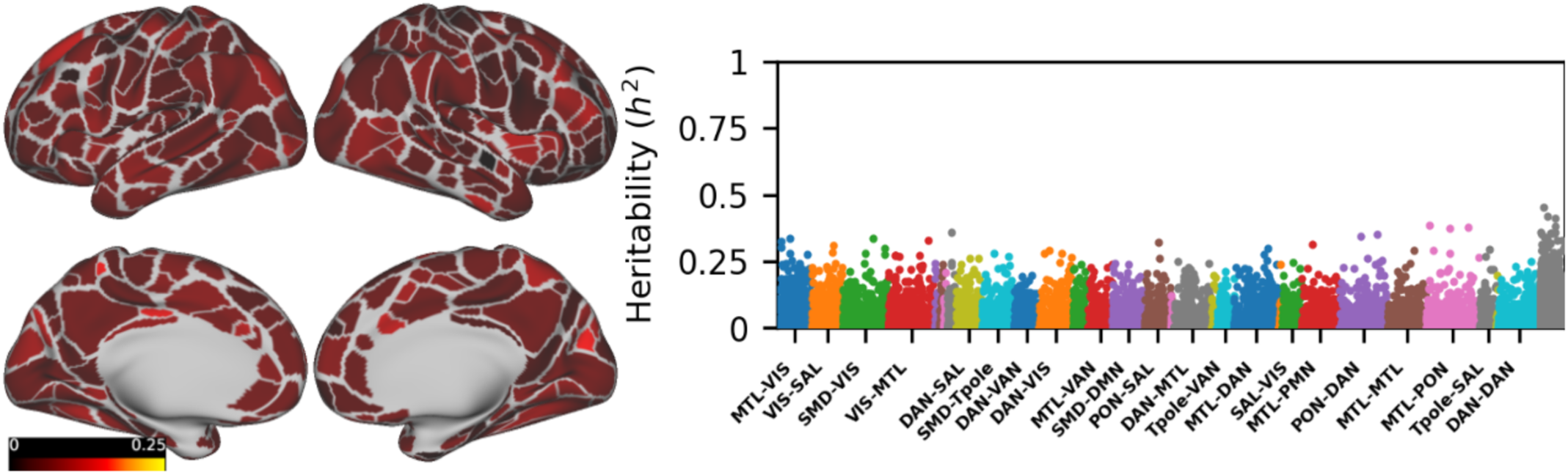
Left: ROI-wise projections of 90th percentile twin heritability estimate for Fisher z-transformed functional connectivity values between Gordon parcels, mapped onto the Conte69 inflated cortical surface. The color scale spans 0 to 25% heritability for consistency across figures. Both lateral (top) and medial (bottom) views are shown for each hemisphere. Right: Manhattan-style plot of heritability estimates, with edges grouped and color-coded by the resting-state networks of the connected ROIs. The y-axis spans the full range of possible heritabilities (0-1). Network-network pairs accounting for fewer than 1.5% of total ROI-ROI connections are rescaled 100-fold on the x-axis and displayed in grey for clarity.

**Figure 4:**
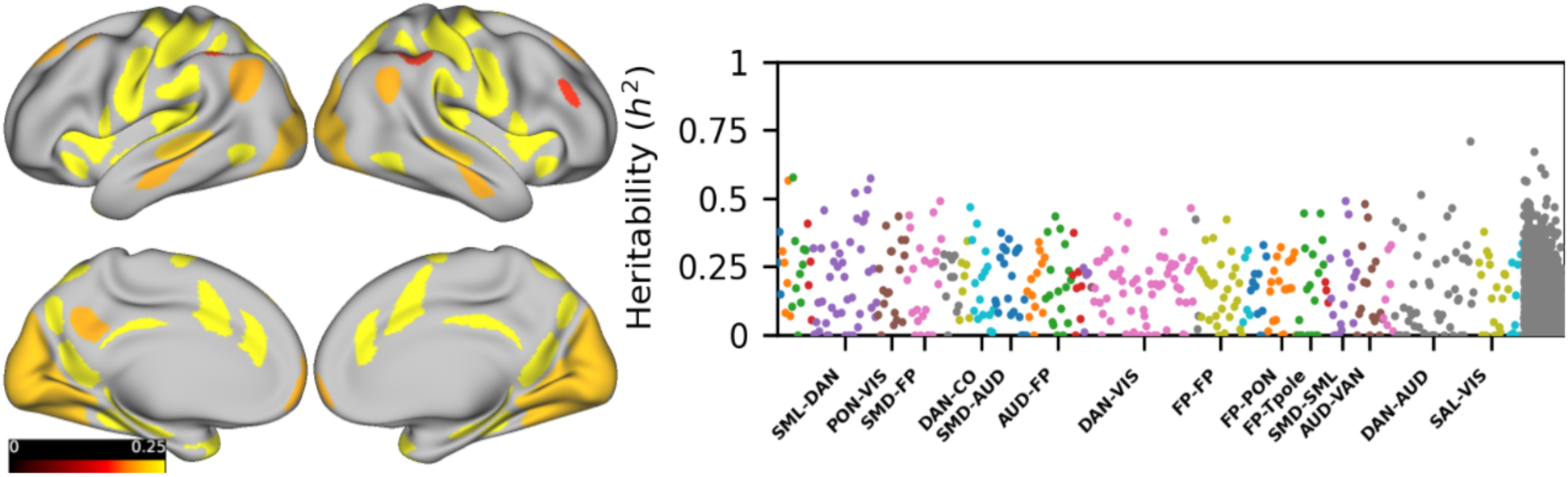
Left: ROI-wise projections of 90th percentile twin heritability estimate for Fisher z-transformed functional connectivity values between MIDB probabilistic parcels, mapped onto the Conte69 inflated cortical surface. The color scale spans 0 to 25% heritability for consistency across figures. Both lateral (top) and medial (bottom) views are shown for each hemisphere. Right: Manhattan-style plot of heritability estimates, with edges grouped and color-coded by the resting-state networks of the connected ROIs. The y-axis spans the full range of possible heritabilities (0-1). Network-network pairs accounting for fewer than 1.5% of total ROI-ROI connections are rescaled 100-fold on the x-axis and displayed in grey for clarity.

SNP and twin-based estimates on FC exhibited poor rank-order concordance and poor consistency in identified significant connections. Kendall’s tau correlations were near zero at both the ROI and network levels (e.g., τ = 0.0015 for Gordon ROIs; τ = –0.03 for probabilistic systems), suggesting that the two methods capture largely non-overlapping sources of variance. For Gordon parcellations, precision and recall of identifying significant connections were 9% and 6%, respectively (F1 = 24%), while for Probabilistic parcellation, precision and recall were 12% and 7% (F1 = 33%).

Additionally, for the estimates on FCs reported here, we observed twin-based heritability estimates, averaging ∼ 33% which are comparable to published twin heritability results (Elliott et al., 2018). This alignment with established values supports the validity of our modeling framework and suggests that the relatively low heritability observed for functional phenotypes does not reflect a general insensitivity of the method but rather points to a more limited genetic contribution to functional brain organization.

### Heritability of functional network topography is small

We next examined individualized functional network topography, defined as the cortical surface area of functionally defined networks. Estimated SNP-based heritability was minimal, with a maximum value of just 2% (**Figure. 5**). Twin-based heritability ranged from 6% to 55%, consistent with moderate twin heritability for a subset of networks. These estimates remained stable after controlling for global cortical surface area (r = 0.99 for twin-based; r = 0.98 for SNP-based estimates). Agreement between twin and SNP identified heritable networks was limited. These findings further suggest either each method is capturing the distinct sources of genetic signal or differences in their respective subsamples.

**Figure 5:**
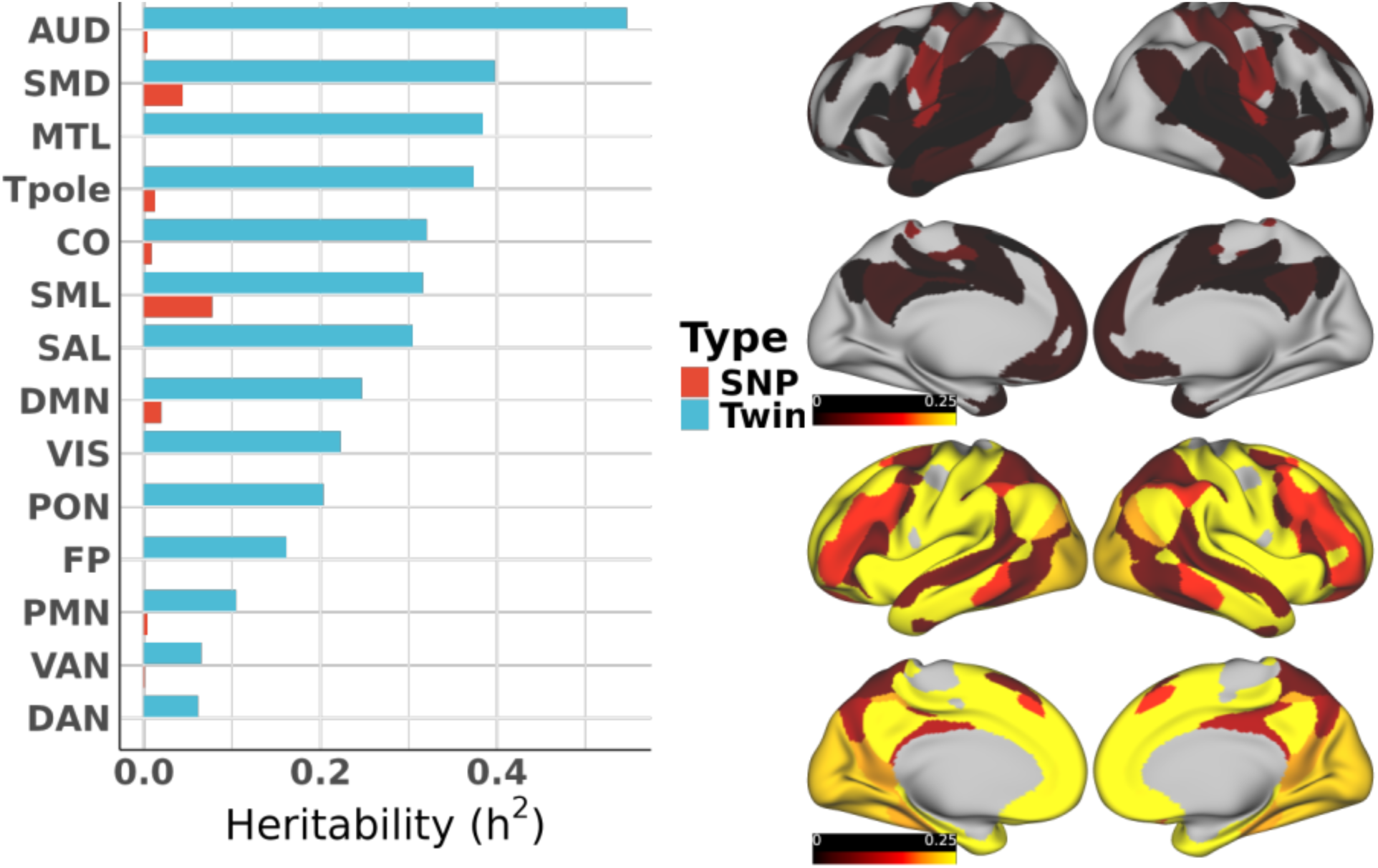
SNP and twin heritability estimates for network surface area as calculated from the PFMs. (left) Results are ordered by the magnitude of the twin heritability estimate along the y-axis. (right) Surface area heritability of each probabilistic functional network is projected onto the surface of the inflated Conte brain for SNP (top) and twin (bottom) estimates. The color scales are restricted from 0-25% heritability for direct comparison. Estimates of exactly zero are omitted.

**Figure 6.**
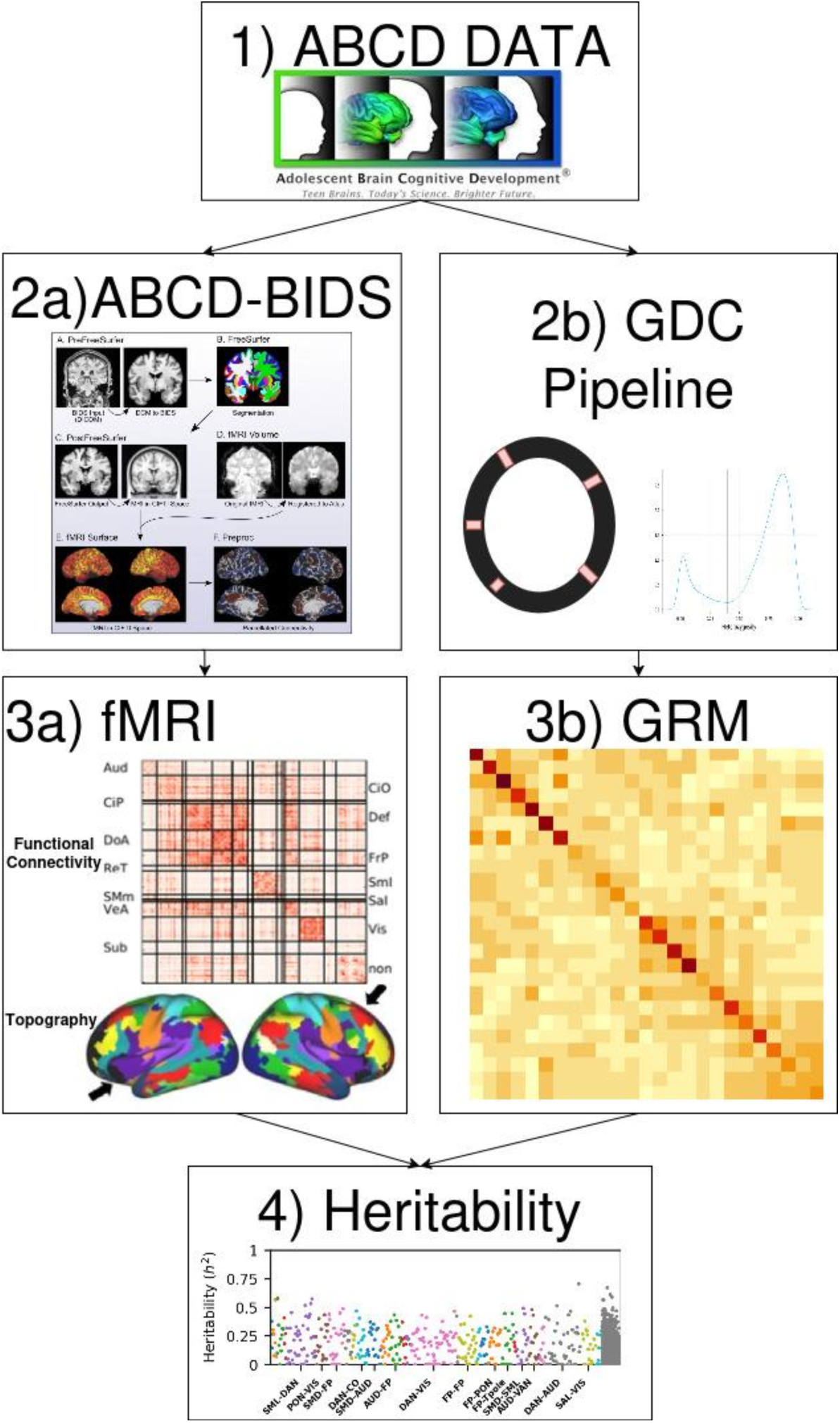
An overview of data pipelines utilized in this study. Imaging data followed the standard ABCD BIDS pipeline (2a) and time series were analyzed to compute pairwise pearson correlations (3a). Smokescreen genetics data were processed using the GDC in-house genomics pipeline (2b) which were then used to compute pairwise genetic correlations, stored in the GRM), between all subjects (3b). The GRM was used, along with standard sociodemographic variables and GRM PC’s as explanatory variables for the imaging derived features to provide heritability estimates using AdjHE-RE (4).

While twin-based models detect modest genetic contributions to network topography, SNP-based heritability estimates reveal that common genetic variation accounts for only a small fraction of this variability. The common environmental effect estimates from twin models were all estimated to be at or near zero with 95% confidence intervals encapsulating zero from every network.

## Discussion

Our study estimated the SNP-based heritability of functional connectivity and network surface area in a large cohort of adolescents (n =11,572, aged 9-10) from the ABCD dataset where we used a subset comprised of 5,247 unrelated individuals, 330 dizygotic, 248 monozygotic twins. We observed that SNP-based heritability estimates for FC derived using group average Gordon parcellations were low (median [5%ile, 95%ile] = 2e-10% [8e-9%, 9%], max = 39%) during adolescence. Similarly, the SNP-based heritability of FC for MIDB probabilistic parcellations, which only consider ROIs that have high spatial consensus between subjects, was low (median [5%ile, 95%ile] = 5.8e-7% [4e-10%, 3%], max = 12%). The heritability for network surface area for individuals was also small with a maximum of 2%. Twin heritability estimates were moderately higher (median [5%ile, 95%ile] = 8.9e-6% [6e-35%, 34%] and 6% [8.5e-31%, 40%] for Gordon and probabilistic parcels and 6% [0%, 39%] for network surface area). Both twin and SNP findings suggest a prominent role for environmental exposures in shaping brain functional organization during or prior to adolescence that agrees with recent findings. Adolescence is a period of rapid brain development (Larsen et al., 2023; Larsen & Luna, 2018), with substantial changes in brain structure and function. Environmental experiences during this time, such as peer interactions (Lecce et al., 2024), family dynamics (Hofstee et al., 2022; Lecce et al., 2024), and nutritional factors (Hofstee et al., 2022; Lecce et al., 2024; Moran & Lowe, 2016), influence brain development. Here we replicate the magnitudes of these estimates to align with previous SNP-heritability estimates found in adults via the UK Biobank and extend those findings to children, which provides additional confidence for our findings reported here. In addition, we replicated heritability estimates found in ABCD (Elliott et al., 2018); Sun et al., 2025) and the relative magnitudes between twin and SNP heritability reflect the “heritability gap” (see below).

The discrepancy between twin heritability and SNP-based heritability of functional brain features observed in our study exemplifies the “missing heritability gap” often discussed in genetics research. While disagreements abound regarding whether twin studies or SNP studies are sufficiently powered (Matthews & Turkheimer, 2022), our findings here both replicate and extend prior literature, therefore we will attempt to synthesize twin and SNP findings here. Twin heritability can be influenced by rare genetic variants, shared environmental factors, or non-additive genetic effects. In particular, twin studies can be subject to genetic confounding, where genetic influences are correlated with environment because twins share the same household. As a result, heritability estimates within most twin studies represent an upper bound on possible heritability (Matthews & Turkheimer, 2022). On the other hand, SNP-based heritability only measures genetic relatedness at the SNP level, ignoring other levels of the genome such as copy number variants or topologically associated domain (TADs). Therefore, SNP-based heritability reflects a lower bound on heritability from additive genetic effects assumed in such models as PRS. While estimating both provides bounds on heritability estimates, measuring SNP-based heritability helps refine inferences for PRS brain correlations.

In summary, our study highlights the importance of considering both genetic and environmental factors when studying the development of functional brain networks and their role in mental health. SNP-based heritability estimates for brain network features were consistently low for adolescents. Combined with prior evidence that SNP-heritability is low for functional connectivity in adults, the findings emphasize the need for careful interpretation of PRS associations with brain function. For example, previous studies have linked PRS derived from Alzheimer’s disease (Axelrud et al., 2019), autism spectrum disorder (Lawrence et al., 2022), ADHD (Hermosillo et al., 2020), general psychopathology (Sun et al., 2025) and schizophrenia (Qi et al., 2022; Cao et al., 2021)) to brain features. The common interpretation of these studies is that PRS associations with brain network features reflect the genetic underpinnings of the function of those networks. However, this interpretation is similar to the sailors example provided in the introduction. Behavioral PRS may be correlated with brain network features because of an indirect trait that increases exposure and alters brain network features as opposed to genetic variants that influence brain network features directly.

While novel and important, our study has several limitations. Our study examines SNP heritability estimates of functional brain network features. However, such an approach only accounts for SNPs that were directly genotyped or imputed with high confidence, which represent only a fraction of the variability observed in the human genome. Non-SNP polymorphisms and epigenetic factors, which also contribute to individual differences, are left unaccounted for. Additionally, while the chosen set of SNPs were sufficient for detecting significant heritability in volumetric measures of the brain for the diverse multisite ABCD study (Coffman et al., 2024), it is possible that heritability of functional brain features are contained in the set of excluded SNPs. Nevertheless, our study is a stepping stone to future work considering the nuances of possibly genetically correlated brain imaging traits. Another limitation is that we are only able to capture a snapshot of genetic influences on functional brain networks due to the cross-sectional nature of the sample. As longitudinal studies like ABCD and Healthy Brain and Child Development (HBCD) study collect more data across time, it will be imperative to examine whether genetic influences to brain network features are stable across lifespan.

## Methods

### ABCD Data and Participants

The Adolescent Brain Cognitive Development (ABCD) study includes N = 11,572 children aged 9–10 years, recruited across 21 sites in the United States (Casey et al., 2018; Uban et al., 2018). The sample is demographically diverse (Garavan et al., 2018), comprising 48% female (range across sites: 43–61%), 52% White (range: 8–87%), 15% Black (range: 0–54%), 2% Asian (range: 0–13%), 20% Hispanic (range: 2.3–74%), and 11% Other (range: 3.6–23%). Participants with missing or incomplete imaging, genomic, or demographic data, as well as those with significant motion artifacts during MRI scans, were excluded from the analysis. The final dataset utilized 5,247 and 578 subjects for the SNP and twin estimates, respectively.

This study used publicly available data from the ABCD consortium. As a secondary analysis of an existing data repository, no additional IRB approval was required. Oversight was jointly governed by the University of Minnesota, the National Institute of Mental Health (NIMH) Data Archive, and the ABCD consortium. A data use agreement was authorized by the University of Minnesota and approved by the NDA.

### Genomic Data

Genomic data were generated using the Smokescreen™ Genotyping Array (Casey et al., 2018; Uban et al., 2018), which includes over 300,000 SNPs. Data were phased and imputed using the Michigan Imputation Server (Das et al., 2016), with automated and manual quality control (QC) steps applied. QC steps included: Variants with minor allele frequency > 5%, variants with Hardy-Weinberg equilibrium p-values > 1 × 10⁻⁶, variants with imputation R² > 0.99, individuals with heterozygosity < 3 standard deviations from the mean heterozygosity, individuals without hidden familial relationships.

After QC, the final genomic dataset included 1,292,075 SNPs. Genetic relatedness matrices (GRMs) were computed using Plink2 to quantify pairwise genetic similarity between participants. The first 30 GRM principal components were derived to account for population stratification and included as covariates in subsequent analyses.

### Image Acquisition, Preprocessing, and Connectivity

Neuroimaging data were acquired using T1-weighted anatomical MRI scans following the ABCD protocol, with a 3D MP-RAGE sequence (1 mm isotropic resolution, TR = 2500 ms, TE = 2.88 ms, TI = 1060 ms, flip angle = 8°) ((Casey et al., 2018; Uban et al., 2018)). Detailed imaging protocols are available at https://abcdstudy.org/images/Protocol_Imaging_Sequences.pdf](https://abcdstudy.org/images/Protocol_Imaging_Sequences.pdf)

All functional MRIs were processed with the publicly available ABCD-BIDS pipeline (https://github.com/DCAN-Labs/abcd-hcp-pipeline), which is a modified version of the Human Connectome Project (HCP) processing pipelines (Feczko et al., 2021). This pipeline was used to minimize artifacts and standardize data preprocessing. The processing pipeline included multiple stages: 1) corrected for MR gradient and bias field distortions 2) PreFreeSurfer brain extraction 3) the DCAN-labs processing pipeline also applies Advanced Normalization Tools (ANTs), including ANTs DenoiseImage to improve structural clarity and ANTs N4BiasFieldCorrection to reduce field bias (Marek et al., 2019).

Parcellation followed the DCAN pipeline, with functional parcellation performed using the Gordon atlas (Gordon et al., 2016) augmented with subcortical volumes defined by FreeSurfer, yielding a total of 352 regions of interest (ROIs). Motion correction was applied during preprocessing, and participants exceeding a predefined framewise displacement threshold of 0.2 mm were excluded. Functional connectivity was computed as the Pearson correlation of quality-controlled (QC’d) time series for each subject, with Fisher z-transformation to transform the results to the real line before further analysis.

### Functional brain measures

Functional connectivity was assessed using the 333 ROIs defined by the Gordon atlas because they were independently identified a priori and validated (Gordon et al., 2016). The Gordon ROI set was augmented by subcortical features defined by FreeSurfer, yielding a total of 352 ROIs total. The preprocessed resting-state fMRI time series were extracted from each ROI by averaging the BOLD signal across all vertices within a parcel. Functional connectivity was then computed by calculating the Pearson correlation coefficient between the mean time series of each pair of ROIs, resulting in a 352 × 352 symmetric connectivity matrix with 61,776 connections in the lower triangle. This matrix represents the strength of temporal co-fluctuations between brain regions, with higher absolute values indicating stronger functional relationships. Fisher’s r-to-z transformation was applied to the correlation coefficients to relate the measures to the real axis and provide more symmetry.

To calculate the functional connectivity of the regions of interest (ROIs), a probabilistic parcellation was derived based on the consistency of network assignments across subjects. A voxel-wise analysis determined the proportion of subjects for which a given grayordinate was assigned to a specific network, creating a probabilistic network map. These maps were thresholded at 75% consensus, meaning that only regions assigned to the same network in at least 75% of subjects were retained which yielded 80 ROIs. The functional connectivity between these thresholded ROIs was then computed, ensuring that only high-consensus regions contributed to the final connectivity estimates. Additionally, clusters smaller than 30 grayordinates were excluded to enhance the reliability of the parcellation leading to 3,600 connections and 3,160 in the lower triangle (Hermosillo et al., 2024).

In order to calculate the surface area of each network, subject-level precision functional network maps were created. Such maps increase signal homogeneity by controlling for individual variability in spatial organization of brain networks. We used template matching procedure (Hermosillo et al., 2024) to create individual network maps for each subject in the dataset. Then, we measured the surface area (in mm²) associated with each vertex on the individual’s mid-thickness surface using Connectome Workbench. Next, we determined the relative contribution (size) of each functional network to the overall cortical surface area by comparing the total surface area of all vertices within a network to the total cortical surface area. Proportional surface area for each network was determined for each individual participant and used for analysis.

### Statistical Analysis

SNP heritability was estimated, which quantifies pairwise genetic similarity while controlling for differences in image acquisition at the site level. Variance components were estimated using the AdjHE-RE method of moments (Coffman et al., 2024) approach on the set of unrelated subjects. AdjHE-RE projects the covariance of the covariate residualized outcome against the genetic relatedness matrix (GRM), a binary matrix specifying whether each pair of subjects was observed at the same site, and the identity matrix. In this way, AdjHE-RE estimates the proportion of variance of the outcome attributable to additive genetic effects as opposed to subject specific variations. The GRM was calculated as the pairwise genetic correlation between all subjects utilizing GCTA (Yang et al., 2011). The first 30 principal components of the GRM were utilized to control for differences in allele frequency from the diverse ancestry in the ABCD dataset. Twin heritability estimates followed the standard ACE model on twin tuples and utilized the mets library (1.3.4) in R (version 4.4.0 (2024-04-24) – “Puppy Cup”) (Eubank & Kupresanin, 2011; Scheike et al., 2014). Both models controlled for age, sex, household income, and scanner site as covariates. All statistical tests were adjusted for multiple comparisons using a Bonferroni correction at α = 0.05 within each estimation-phenotype set. Corrections were applied independently for each set, instead of accounting for the total number of tests across different estimation-phenotype sets. Estimates were constrained to be nonnegative. Estimates were generally summarized by the resulting heritability estimate and sets of estimates of estimates were summarized by their central fifth and ninety fifth percentiles except in cases where they were exceedingly close to zero when they are simply summarized as the maximum estimate within the set. Statistical visualizations were made in ggplot2 (3.5.1) and brain visualizations were made in Connectome workbench (2.0.0). Data storage and computing were done using supercomputers at the Minnesota Supercomputing Institute (MSI) at the University of Minnesota.

## Notes

### Competing Interest Statement

The authors have declared no competing interest.

